# Neuronal contact predicts connectivity in the *C. elegans* brain

**DOI:** 10.1101/2022.11.13.516316

**Authors:** Steven J. Cook, Cristine A. Kalinski, Oliver Hobert

## Abstract

Axons must project to particular brain regions, contact adjacent neurons, and choose appropriate synaptic targets to form a nervous system. Multiple mechanisms have been proposed to explain synaptic partnership choice. In a ‘lock-and-key’ mechanism, first proposed by Sperry’s chemoaffinity model^1^, a neuron selectively chooses a synaptic partner among several different, adjacent target cells, based on a specific molecular recognition code^2^. Alternatively, Peters’ rule posits that neurons indiscriminately form connections with other neuron types in their proximity; hence, neighborhood choice, dictated by initial neuronal process outgrowth and position, is the sole predictor of connectivity^3,4^. However, whether Peters’ rule plays an important role in synaptic wiring remains unresolved^5^. To assess the nanoscale relationship between neuronal adjacency and connectivity, we evaluate the expansive set of *C. elegans* connectomes. We find that synaptic connectivity can be accurately modeled as a path-length-dependent process of neuronal adjacency and brain strata, offering strong support for Peters’ rule as an organizational principle of *C. elegans* brain wiring.

## Results and Discussion

The main component of the *C. elegans* brain is its nerve ring, a bundle of neurites with extensive *en passant* synapses (Fig 1). Nearly all neurons present in the nerve ring exhibit axonal characteristics and connect to a subset of their neighbors. Neurites projecting into the nerve ring cluster into multiple groups/strata with similar functions based solely upon the patterns by which they are physically adjacent^6,7^ (Fig 1C). When visualized together from volumetric serial section EM (ssEM) reconstructions, these neurite bundles minimally overlap with other bundles or strata^6,7^. We refer to these bundles throughout as ‘strata classifications’, whose geometric abstraction imposes spatial constraints on the *C. elegans* brain.

**Figure 1.**
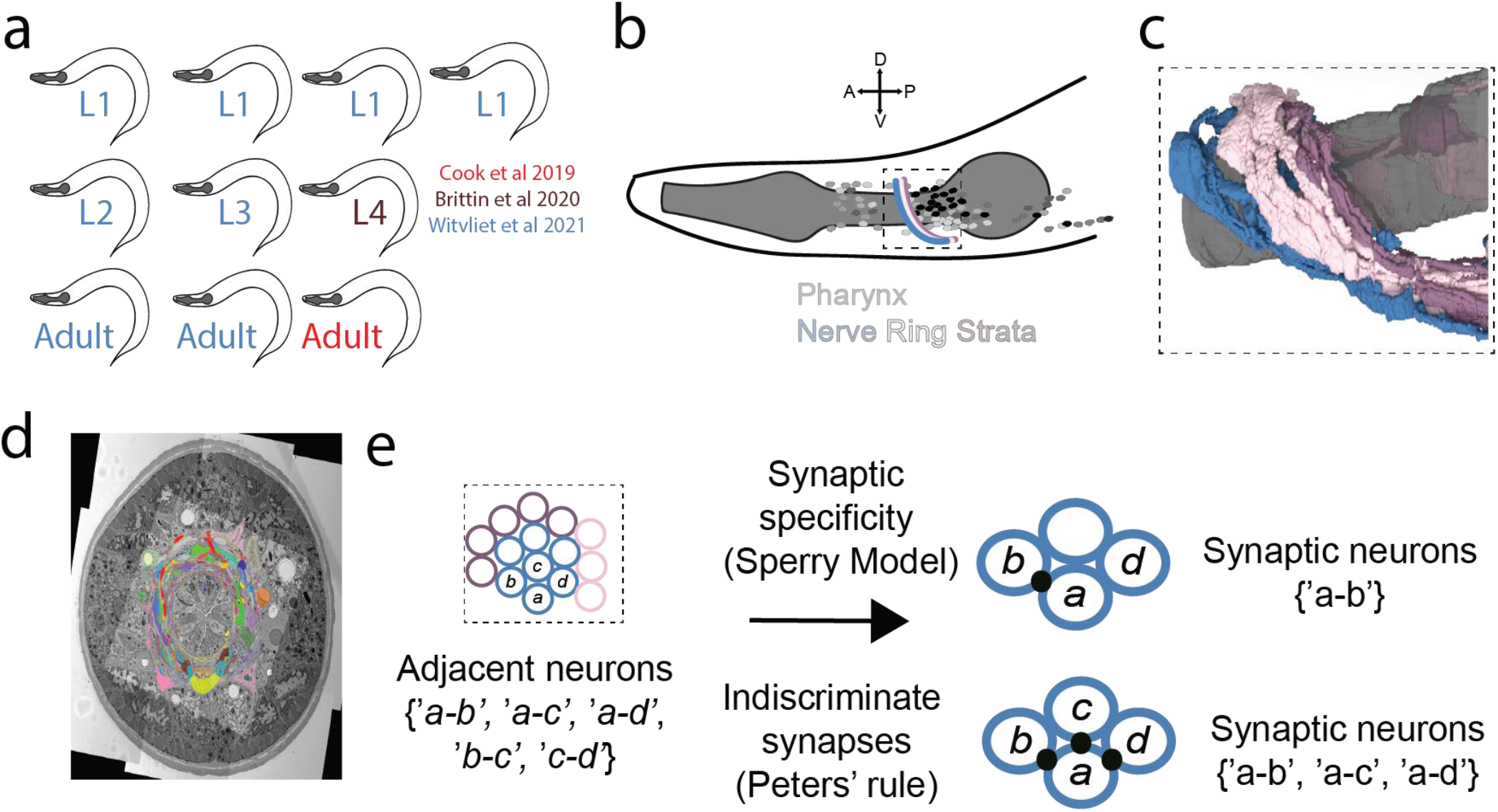
Structure of the *C. elegans* nervous system and models of synaptic specificity. A) Graphical depiction of animals sampled for this study. Animals from Cook et al 2019 are in red, Brittin et al 2020 in brown, and Witvliet et al 2021 in blue. B) Cartoon of the *C. elegans* anterior, with the region of our study presented in a dashed-line box, the nerve ring is multi-colored, neuronal cell bodies are small, shaded ovals, and the pharynx is in dark gray. C) Lateral view of 3D reconstruction showing the pharyngeal isthmus (gray) and three nerve ring strata (blue, pink, purple). D) Example electron micrograph showing a dense reconstruction of the nematode nerve ring. E) Simple depiction of how synaptic specificity may be achieved between adjacent neurons. Either a neuron may selectively choose among adjacent partners, or indiscriminately synapse with all adjacent partners.

How is wiring being specified in the *C. elegans* brain, and does neuronal contact dictate connectivity? White first estimated that neurons make synaptic connections with 45% of their adjacent neighbors^8^. From an analysis of legacy connectomic data, it was concluded that the probability of synapse formation is practically independent of the weight of adjacency between two processes^9^. These groundwork analyses suggested that Peters’ rule may not be an organizing principle of *C. elegans* brain wiring.

Several recently published wiring diagrams of the *C. elegans* brain^6,10^ allowed us to re-consider the applicability of Peters’ Rule as an organizational principle of *C. elegans* brain wiring. In particular, the availability of multiple, brain-wide wiring diagrams that capture both synaptic connections (“connectome”) and cell surface adjacencies (“contactome”) allowed us to take novel perspectives on variability and stereotypy of the connectome as compared to single or few sample studies. Together, these datasets represent the largest and most complete sampling of any animal’s wiring diagram to date (Supplemental data 1). To create a composite dataset that accurately compares differences within and across individual EM samples, we made several assumptions and only evaluated embryonically born neurons by cell class^11^. For full assumptions, see details in the methods section. Our combined dataset of 2683 adjacencies and 1404 connections represents our goal of making unbiased comparisons between wiring diagrams.

Despite its compact and conserved structure, the *C. elegans* nervous system is highly interconnected. Calculating from our aggregate dataset, neurons in the *C. elegans* brain have a high probability of encountering each other throughout development. On average, a neuron contacts 79.5% of all other neurons across samples (Fig 2A). Neurons are more selective with their choice of synaptic partners, making connections with an average of 38.6% of all other neurons (Fig 2A). We next calculated the filling fraction or proportion of adjacencies that also have a synaptic connection for each sample. The aggregate filling fraction was 0.523, while the mean of individual filling fractions was 0.303 (Fig 2B). These data update and increase previous estimates of filling fraction^8,10^, further exhibiting that *C. elegans* has a wide range of possible neuronal adjacency and connectivity configurations.

**Figure 2.**
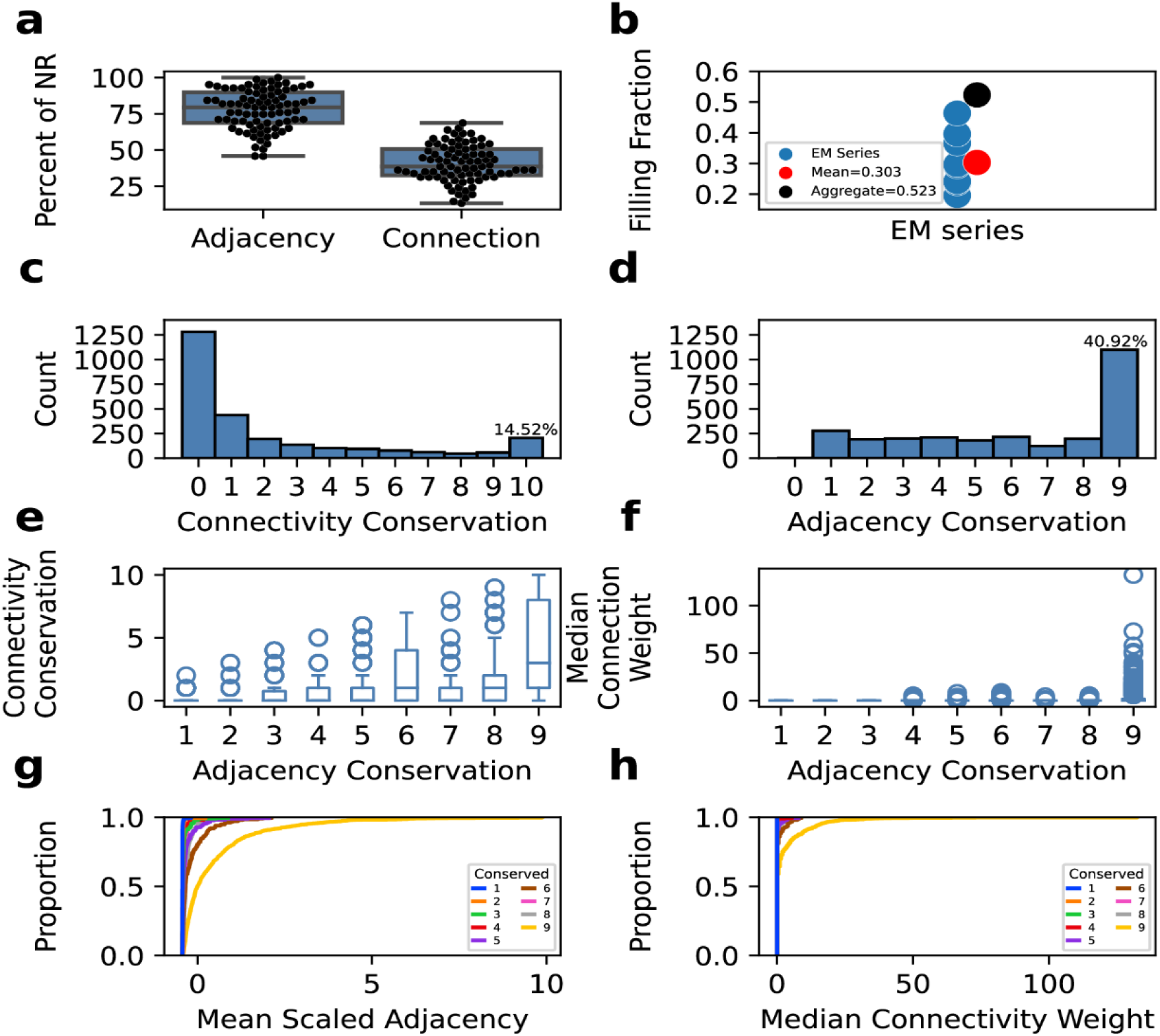
Presence and sources of variability in the *C. elegans* brain. A) Boxplots of percent adjacency and connections by neuron class. B) Dot plot of filling fraction, or percent of adjacencies that result in a connection, of each EM series evaluated with mean and aggregate in red and black, respectively. C) Histogram of connectivity conservation across EM series. 14.52% of undirected connections are found in each EM series. D) Histogram of adjacency conservation across EM series. 40.92% of adjacencies are found in each EM series. E) Boxplot of adjacency conservation vs synaptic conservation. F) Boxplot of adjacency conservation vs median connection weight. G) Cumulative distributions of mean scaled adjacency vs proportion of adjacencies, colored by conservation across EM series. H) Cumulative distributions of median connectivity weight vs proportion of connections, colored by conservation across EM series.

How repeatable are adjacencies and connectivity across individual samples? We calculated a conservation score for each connection and adjacency in our dataset, representing the number of samples where that edge is present. The most frequently observed connectivity conservation level is 0, where two neurons are adjacent but asynaptic (Fig 2C). The second-most observed conservation is 1, indicating our sensitivity to ‘rare’ synaptic connections. 14.52% of connections are observed across all EM samples. We next asked whether variability in connectivity could be largely driven by differences in adjacency. The distribution of adjacency is different from connectivity, where the most common observed adjacency is completely conserved across samples (Fig 2D). These distributions have two implications: First, adjacency is more conserved than connectivity, and, second, the existence of a ‘core circuit’, indicated by a previous analysis^6^, is confirmed by a larger sample size.

Because both neuronal adjacency and connectivity are variable, we asked whether these two metrics share any commonalities. We plotted connectivity conservation against adjacency conservation and found that while adjacency conservation can set an upper bound for connectivity, it does not strongly relate to connectivity conservation. (Fig 2E). We plotted our weight metric for synaptic connections against adjacency conservation and similarly observed that completely conserved adjacencies display the widest variability in median connection weight (Fig 2F). The influence of completely conserved adjacency on edge size is also exhibited when plotted as a cumulative density function of the mean scaled adjacency (Fig 2G) or median connection weight (Fig 2H). Together, this suggests that adjacency is more conserved than connectivity across individuals and that completely conserved adjacencies can yield many different configurations (connection conservation) as well as weights (connection size). The extent of variability across individuals disagrees with a generalizable, selective ‘lock-and-key’ mechanism of connectivity in the *C. elegans* brain.

To assess whether a temporal order of synapse and adjacency formation could predict connectomic properties, we evaluated the developmental trajectory of the connectome. Plotting our filling fraction as Witvliet et al 2021, we also find an increase in filling fraction as the animal passes through postembryonic developmental stages, which confirms that many adjacencies precede connections (Fig 2 supplement 1A). We next asked whether synaptic connections could precede a strengthening of adjacencies, i.e., strong connections zipper together adjacent processes. By plotting the adjacencies of completely conserved (Fig 2 supplement 1B), incompletely conserved (Fig 3 supplement 1C), and absent (Fig 3 supplement 1D) connections across development, we find no relationship between connection conservation and scaled adjacency. These results, while correlative, argue against a model where neuronal adjacencies are a consequence of synapse formation.

If the properties of the connectome across individuals or development do not predict a direct relationship between adjacency and connectivity, could the quantitative extent of adjacency instead instruct connectivity? To visualize a possible path-length-dependent relationship between adjacency and connectivity, we plotted mean scaled adjacency against median connection weight. We observed a moderate linear correlation (Pearson’s r=0.682) (Fig 3A) between these metrics; however, many neuronal adjacencies do not have a corresponding synaptic connection (Fig 2C, Fig 3A). Two possibilities for asynaptic adjacencies are large differences in neurite fasciculation to different strata as well as small differences in axonal fasciculation across individuals.

**Figure 3.**
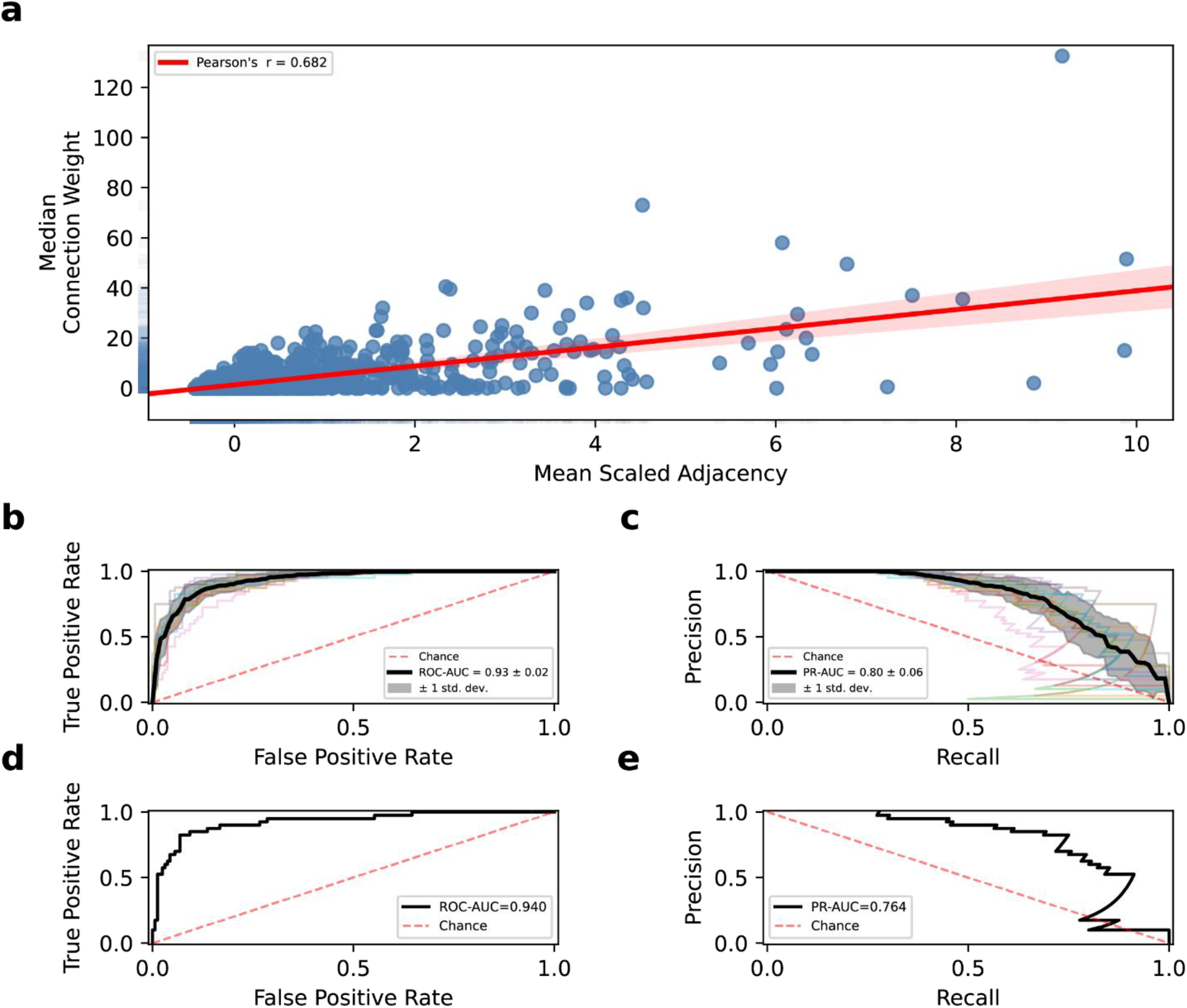
Modeling adjacency as a predictor of connectivity. A) Blue scatter and rug plots of mean scaled adjacency vs median connection weight with linear regression line shown in red, Pearson’s r=0.682. B) ROC curves of the cross-validated model (kfolds=10), black line is the mean ROC-AUC=0.93 +/ 0.02. D) PR curves of the cross-validated model (kfolds=10), black line is the mean PR-AUC=0.80+/0.06. D) ROC curve of final model performance, ROC-AUC=0.95 E) PR curve of final model performance, PR-AUC=0.764.

To address these two possibilities, we applied a binary code for whether two neurons are in the same or different strata. We then used this strata classifier and scaled adjacency and connection weights found in the individual samples to determine their correlation across samples. While strata are correlated with both adjacency and connectivity, we found a consistently stronger relationship between individuals (Fig 3 supplement 1 A,B). Therefore, a quantitative comparison of adjacency and connectivity appeared appropriate.

To reflect the underlying connectomic properties, we modeled synaptic connectivity as a consequence of neurons being directed to a specific brain region and the extent of their adjacency to their neighbors. Our machine learning-based classification problem uses a composite dataset that includes all adjacencies and connections found across animals and includes three predictors and one target variable. Dataset and full model selection criteria are detailed in the methods section. In brief, the predictors are mean scaled adjacency and strata classification (same or different)^6^. The target variable was a binarized connectivity value of median connection weight > 0 vs median connection weight = 0. We adjusted for imbalance in our classifier and reported multiple performance metrics for each model. After splitting the data into a training and test set, we cross-validated the performance of multiple classifiers on our training set (Table supplemental 1). Although the classifiers performed similarly, the logistic regression classifier was chosen for its performance, interpretability, and simplicity (Fig 3B,C). We named this model Nematode Connectivity Classification Model (NCCM) and refer to it by this name hereafter. Because of our imbalanced synaptic classifier, we do not rely on accuracy alone to assess our model. Applying NCCM to the test set yields ROC-AUC (ability to discriminate between positive class and negative class), PR-AUC (precision score per recall threshold), and weighted F1 (harmonic mean of precision and recall) scores of 0.940, 0.76, and 0.88 respectively (see Methods) (Fig 3D,E). NCCM is more capable of predicting connectivity overall than a simple linear correlation alone (Fig 3A). This suggests that NCCM is both precise and robust at predicting connectivity from adjacency and brain strata information. Acutely, our logistic regression model approximates a path-length-dependent threshold of physical contact which must be met to yield a synapse between neurons. We conclude that the probability of two neurons being connected is primarily driven by the amount of neuronal adjacency between pairs of neurons and secondarily by their brain strata.

Lastly, we tested NCCM’s predictive ability as a path-length-dependent model in a different region of the *C. elegans* nervous system. Separated from its brain, *C. elegans* has a small, self-contained pharyngeal, or enteric^12^, circuit which directly connects to its brain through a single neuron (Fig 4A). The pharyngeal nervous system is composed of 20 neurons that fall into 14 classes, making several hundred synaptic connections.

**Figure 4.**
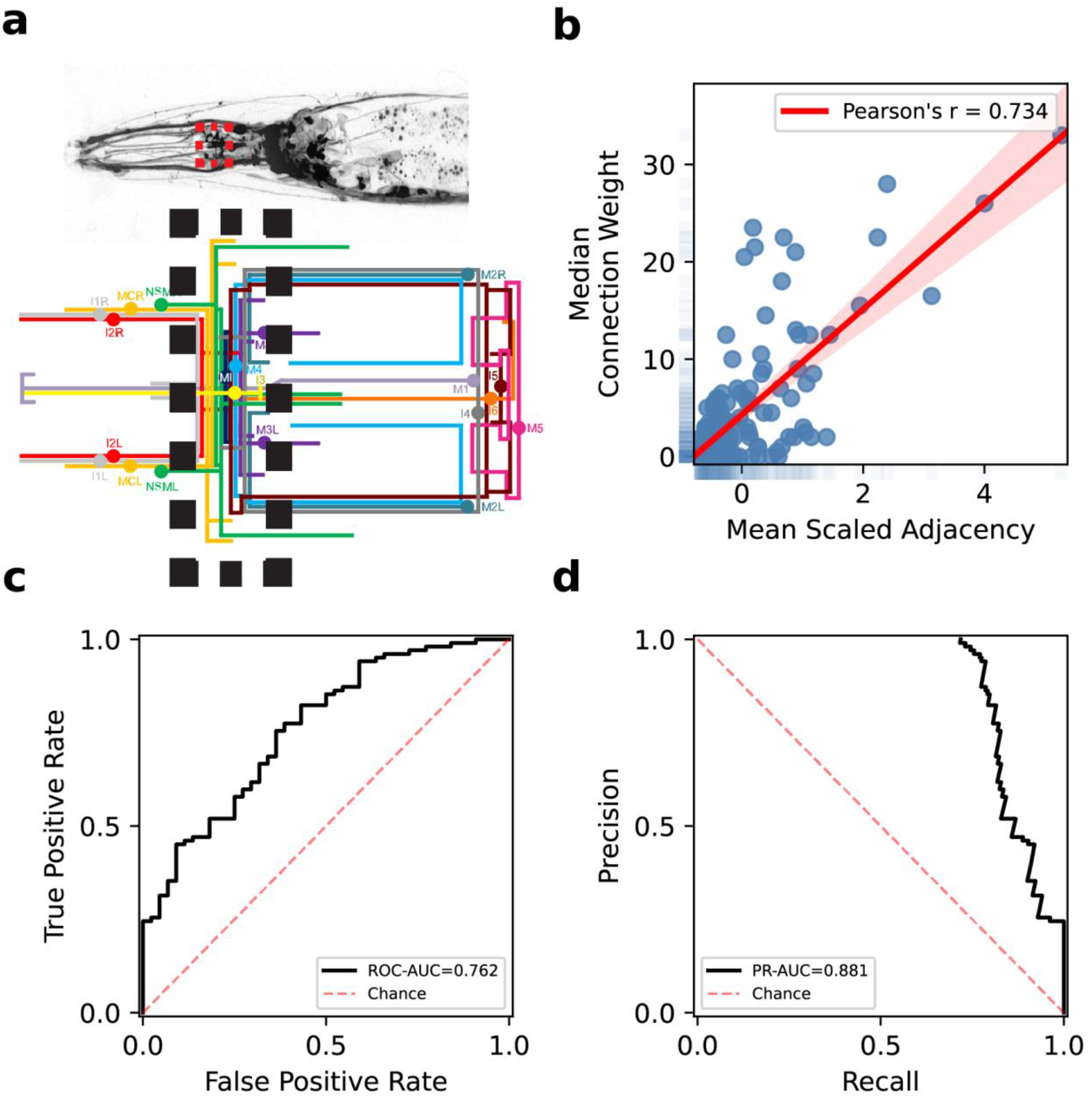
NCCM performance extends to the pharyngeal nervous system. A) (top) Micrograph of all neurons in the *C. elegans* anterior with the pharyngeal nerve ring outlined by a red dashed box subway line depiction of the pharyngeal nervous system with the pharyngeal nerve ring highlighted in a black dashed box B) Blue scatter and rug plots of mean scaled adjacency vs median connection weight with linear regression line shown in red, Pearson’s r= 0.734. C) ROC curve of NCCM performance, ROC-AUC=0.762 D) PR curve of NCCM performance, PR-AUC=0.881.

Previous work showed that the pharyngeal nervous system is more promiscuous than its somatic counterpart, i.e. its filling fraction is larger^13^. We applied NCCM to the densest region of the pharyngeal nervous system, treating the pharyngeal nerve ring (n=2) as an unstratified structure (Supplemental dataset 2). Similar to the somatic nerve ring, we also found a moderately strong correlation between our adjacency and connectivity weights in the pharynx, despite the abundance of neuronal adjacencies without a connection (Fig 4B). The performance criteria for our pharyngeal NCCM were strong, with ROC-AUC, PR-AUC, and F1 scores of 0.762, 0.881, and 0.58, respectively. These results suggest that NCCM was more precise but less robust when applied to the pharynx compared to the somatic nerve ring. Together, our results suggest that NCCM is robust to different brain regions, and different regions of the *C. elegans* nervous system may share fundamental rules of synaptogenesis.

## Conclusions

Here, we provide evidence for neuronal adjacency predicting chemical synaptic connectivity in a path-length-dependent model. Despite extensive inter-individual variability in the contactome and connectome of *C. elegans*, we were able to discern constancies and model synaptic specificity by leveraging an aggregate dataset of recently published wiring diagrams^6,10,13,14^. Our results build upon several recent efforts and represent a new interpretation of how nematode wiring specificity may be achieved. Firstly, our model is explanatory: a neuron is directed to specific brain strata after which a connection is formed if a threshold of physical contact is met. Secondly, our model is comprehensive: while we are not the first study to show the importance of neuronal adjacency in predicting connectivity^6^, we use all available datasets and all observed adjacencies and synapses to capture a new and holistic view of chemical synaptic connectivity in the *C. elegans* brain. Lastly, our model is consistent with decades of *C. elegans* neurogenetic research. Despite extensive genetic screens searching for molecules that confer synaptic specificity, few molecules have been described that operate as keys-and-locks to select synaptic partner choice, as predicted by Sperry. Contrastingly, the pursuit of axon guidance and fasciculation molecules that determine neighborhood choice (i.e. adjacency) during axon outgrowth has been more productive^15,16^.

In conclusion, in *C. elegans*, synaptic target choice appears to be driven by relative axodendritic placement and, hence, initial axodendritic outgrowth and selective fasciculation are the primary determinants of synaptic connectivity. Like the genomics field, we argue that new wiring diagrams will increase our appreciation of dynamics, inter-individual variability, and population differences. Despite variability, we show that both constant and variable features of the nervous system can be reconciled together to address principles of circuit formation. New replicate wiring diagrams of more complex brain structures will be needed to assess Peters’ rule as a general mechanism for nanoscale wiring. It is possible that as brains evolved to have more neurons, more distinct neuron types, and more elaborate structures there was a commensurate increase in the ‘rulemaking’ of synaptic connectivity (e.g. molecular code).

## Acknowledgments

We thank Daniel J. Bumbarger, David Miller III, John G. White, Eviatar Yemini, Kang Shen, Emily Bayer, and Cindy Reyes for the discussion of this manuscript. This work was funded by the Howard Hughes Medical Institute.

## Author contributions

SJC – conceptualization, methodology, investigation, formal analysis, visualization, editing, writing

CAK – investigation, formal analysis, visualization, editing, writing OH – editing, writing, funding acquisition

## Methods

### Data sources

Nerve ring connectivity data are derived from^14^, and^10^. Nerve ring adjacency data for nerve ring data are derived and^10^. Neuronal strata classifications are derived from Brittin et al 2021^6^ and Moyle et al 2021. Pharyngeal connectivity and adjacency data are from Cook et al 2020 ^13^.

### Code availability

All code and data used to perform this analysis and generate figures can be found at https://github.com/stevenjcook/adjacency_connectivity.

### Experiment Methodology

Data preprocessing methods and classification algorithms were performed using Python, including the scikit-learn machine learning library^17^, and the XGBoost gradient boosting library^18^.

### Modeling Assumptions

We evaluated only embryonically-born and sex-shared (i.e. present in both sexes) neurons. The majority of *C. elegans* neurons are left-right symmetric, and we, therefore, grouped and aggregated data by neuronal class (nodes). Adjacencies and chemical synaptic connections (edges) were treated as undirected edges unless noted otherwise. To adjust for size differences across individuals, we scaled individual adjacencies within each dataset. As a proxy for synaptic strength, we counted the total number of individual synaptic connections between neurons. Gap junctions are excluded from our analysis due to their ambiguous ultrastructural characteristics.

### Data Preprocessing

We limited our analysis to adjacencies and chemical synapses between neuronal classes (as compared to individual neurons) that innervate the NR embryonically. Adjacency edge weights are undirected sums of individually adjacent membrane profiles^6^. Connection edge weights are undirected reciprocal sums of connectivity. To account for the differences in adjacency sizes and scoring across samples, we scaled the 9 samples with an available volumetric reconstruction individually using a standard scaler which removed the mean and scaled to unit variance. We then created a new feature, ‘ave_scaled_adjacency’ by taking the mean of the scaled adjacencies across these 9 series. This permits future implementation of our model irrespective of the volumetric reconstruction sample size. For each neuron, we used the strata classification as defined by^6^ and^7^. We created boolean variables to assign whether each row had neurons in the same or different strata (brittin/moyle_bool_1 and brittin/moyle_bool_2). We generated our target variable (dummy_size) by taking the median of the connection weights across all 10 samples, and subsequently binarizing this value (>0 connection weight = 1, 0 connection weight = 0).

### Experimental Setup

The dataset used in our model (NR-modeling.csv) uses four features with ‘ave_scaled_adjacency’, ‘brittin_bool_1’, and ‘brittin_bool_2’ used as predictors and ‘dummy_size’ used as the target variable. The data were split randomly into either the training set or the test set, where 75% of the instances were placed in the training set, and 25% were placed in the test set. The training set was further split into a second training set and a validation set, where 75% of the instances were placed in the training set, and 25% were placed in the validation set. To address the class imbalance in our data, we used Synthetic Minority Over-sampling Technique for Nominal and Continuous (SMOTENC) on our training set^19^.

### Classification Algorithms

The algorithms we evaluated were MLPClassifier, DecisionTreeClassifier, RandomForestClassifier, and LogisticRegression from the scikit-learn library^17^, as well as the XGBClassifier from the XGBoost library^18^.

### Classifier Evaluation

To evaluate the performance of the classifiers, we used a stratified k-fold cross-validation (k=10) and took the mean of a variety of metrics including area under the curve (AUC) the receiver operating characteristic (ROC) curve (ROC-AUC), AUC of the Precision-Recall (PR) Curve (PR-AUC), and F1 score. We used these metrics because we were concerned about false positives arising from an imbalanced target variable. We tested performance of the classifiers using strata classifications defined by either ^6^ or ^7^ to see which resulted in a feature of higher relative importance.

### Model Selection and Performance

Because the classifiers performed similarly across the three metrics, we chose to use logistic regression for our model due to its interpretability. We used a grid search to optimize the parameters of the logistic regression model. The Brittin et al. 2021 classifications were used due to their higher relative importance in the classifiers. We used the training set from the original training and test split, performed SMOTENC, and fit the model.

### Analysis of the pharyngeal nervous system

We further tested our model using an identical training set but with the pharyngeal dataset as the test set. The pharyngeal dataset ^13^(pharynx_modeling.csv) was created the same way as the somatic nerve ring dataset except that only two samples were used, and all neurons were automatically classified as being in the same strata.

## Supplemental figures

**Figure 2 Supplemental 1:**
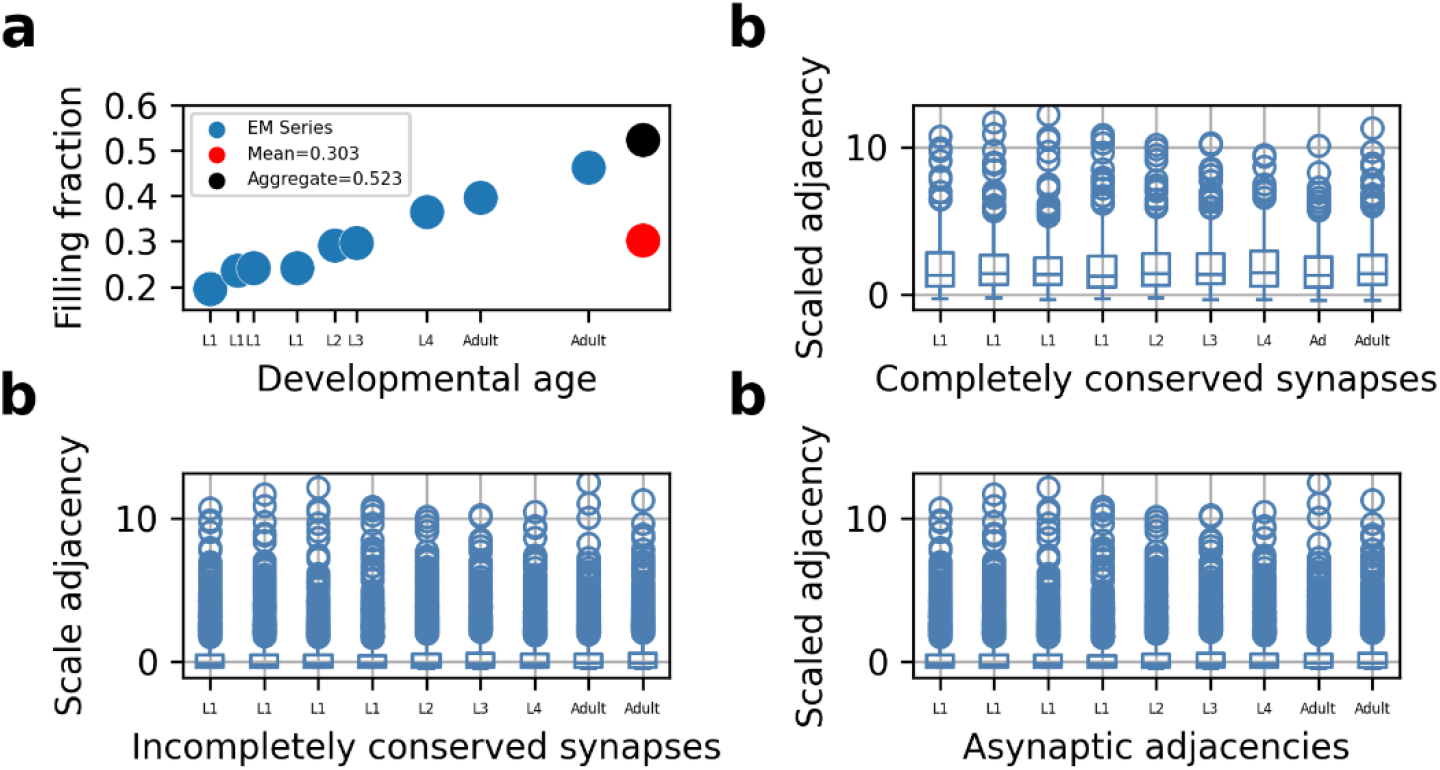
**A)** Dot plot of filling fraction (as seen in Fig 2B) by developmental age. B) Boxplot of scaled adjacency values by developmental age for connections that are completely conserved across samples. C) Boxplot of scaled adjacency values by developmental age for connections that are incompletely conserved across samples. D) Boxplot of scaled adjacency values by developmental age for asynaptic but adjacent neuron pairs across samples.

**Figure 3 supplemental 1:**
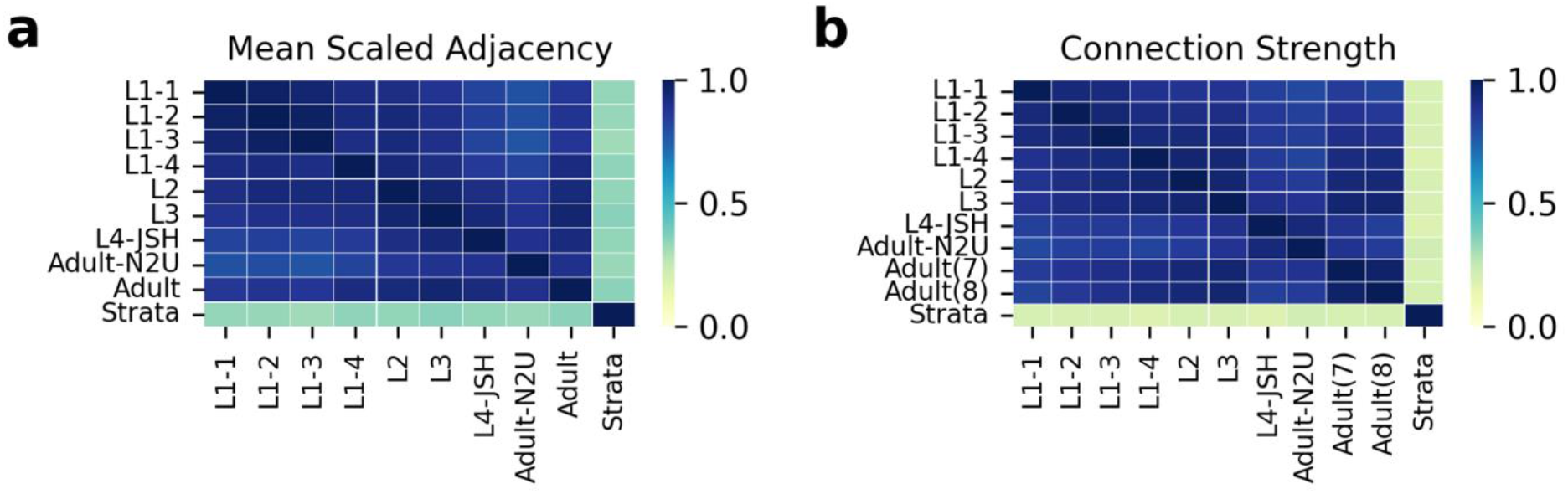
A) Heatmap of Pearson’s correlations between scaled adjacency values of all samples and the binarized strata value. B) Heatmap of Pearson’s correlations between scaled connection weights of all samples and the binarized strata value

**Table supplemental 1.** Performance of multiple ML models. ROC_AUC, PR_AUC, and F1 are reported for all models tested.

| Metric | Multi-layer<br>Perceptron | Random<br>Forest | XGBoost | Decision<br>Tree | Logistic<br>Regression |
| --- | --- | --- | --- | --- | --- |
| ROC_AUC | 0.937600475 | 0.90471884 | 0.915540 | 0.9239594 | 0.93730694 |
| PR_AUC | 0.791002832 | 0.69831236 | 0.653441 | 0.7032076 | 0.79090089 |
| F1 | 0.706096694 | 0.67746561 | 0.537985 | 0.7093193 | 0.730452724 |

**Supplemental data 1: nerve ring modeling**. Aggregate dataset of adjacency and connectivity of all datasets.

**Supplemental data 2: Pharyngeal modeling**. Aggregate dataset of adjacency and connectivity for the pharyngeal datasets.

